# Machine learning prediction of future amyloid beta positivity in amyloid-negative individuals

**DOI:** 10.1101/2023.07.17.547202

**Authors:** Elaheh Moradi, Mithilesh Prakash, Anette Hall, Alina Solomon, Bryan Strange, Jussi Tohka, for the Alzheimers Disease Neuroimaging Initiative

## Abstract

**INTRODUCTION:** The pathophysiology of Alzheimer’s disease (AD) involves *β*-amyloid (A*β*) accumulation. Early identification of individuals with abnormal *β*-amyloid levels is crucial, but A*β* quantification with positron emission tomography (PET) and cerebrospinal fluid (CSF) is invasive and expensive.

**METHODS:** We propose a machine learning framework using standard non-invasive (MRI, demographics, APOE, neuropsychology) measures to predict future A*β*-positivity in A*β*-negative individuals. We separately study A*β*-positivity defined by PET and CSF. RESULTS: Cross-validated AUC for 4-year A*β* conversion prediction was 0.78 for the CSF-based and 0.68 for the PET-based A*β* definitions. Although not trained for the clinical status-change prediction, the CSF-based model excelled in predicting future mild cognitive impairment (MCI)/dementia conversion in cognitively normal/MCI individuals (AUCs, respectively, 0.76 and 0.89 with a separate dataset).

**DISCUSSION:** Standard measures have potential in detecting future A*β*-positivity and assessing conversion risk, even in cognitively normal individuals. The CSF-based definition led to better predictions than the PET-based definition.

## INTRODUCTION

Alzheimer’s disease (AD) is a common neurodegenerative disease with a complex and unclear pathway and a long prodromal phase. The progressive and irreversible nature of AD highlights the need for detecting early changes in the brain that occur decades before dementia. Research on amyloid, tau, and neurodegeneration (ATN) biomarkers, in accordance with the NIA-AA 2018 framework ^1^, has focused on tracking AD progression through beta-amyloid (A*β*) and Tau protein accumulations. These biomarkers are associated with neurodegeneration and cognitive decline ^2^. Identifying amyloid burden in cognitively normal individuals holds promise for identifying those at risk of developing AD ^3^, and it is expected to become the standard for prescribing A*β*-targeted drugs ^4^. Currently, cerebrospinal fluid (CSF) and positron emission tomography (PET) with 18F-labeled amyloid tracers are established methods for confirming the presence and measuring the extent of A*β* accumulation in the brain ^5^. PET can detect metabolic and biochemical alterations in the brain deviating from normal. The clearance efficiency of A*β* protein aggregates can be detected in CSF ^6^. CSF peptides (A*β*_1-42_) and hyperphosphorylated tau are correlated with amyloid plaques and neuronal tangles observed in brain autopsies ^7^ and are linked to cognitive decline, offering valuable insights into early detection. However, PET and CSF are not widely available. CSF can cause discomfort and are invasive, while PET involves exposure to radiation and requires specialized equipment and personnel. Furthermore, there is an ongoing debate over the accuracy of A*β*-positivity determination based on cut-off values from these methods and their mutual concordance ^8,9^. Therefore, promoting the use of universally available data, such as demographics and cognitive scores, is important to advance AD research ^10,11^.

Standardized and longitudinal datasets such as ADNI (Alzheimer’s Disease Neuroimaging Initiative), provide valuable resources for developing machine learning (ML) models with different feature combinations to study PET and CSF biomarkers. These ML models can highlight the important modalities for identifying at-risk individuals transitioning from normal control (NC) to mild cognitive impairment (MCI) and also potentially to dementia by tracking changes in A*β*-positivity states. However, there are limited studies addressing the future predictability of A*β*-positivity using widely available measures ^12^. Additionally, the comparative analysis of model performance based on categorizing individuals as A*β*-positive or A*β*-negative using either CSF or PET biomarkers remains understudied.

This study aims to develop an ML-based approach to predict the conversion to A*β*-positivity in individuals who are A*β*-negative using widely available data in clinical settings. We utilize demographic data (age, gender, education), APOE4 (genetic), neuropsychological scores, and MRI-derived brain volumes to predict whether an individual with A*β*-negative will convert to A*β*-positive (referred to as Ab-Converter) or remain A*β*-negative (referred to as Ab-Stable) over a 4-year period. Analyses are performed separately and in parallel for grouping individuals based on CSF and PET biomarkers. We analyze the role of different data types in the predictive performance of the model in each CSF and PET-based cohort. Additionally, we examine whether baseline CSF/PET measurements improve the predictive performance of the model and how that affects the contribution of cognitive measures and MRI biomarkers. We also investigate the role of baseline PET measurements in predicting A*β*-positivity based on CSF (A*β*42) and vice versa, the role of baseline CSF measurements in predicting A*β*-positivity based on PET. Finally, we evaluate our model’s predictive capability for conversion to MCI/dementia in healthy and MCI individuals. Our study extends to demonstrate that both classification and regression approaches display similar trends.

## MATERIALS AND METHODS

### ADNI Data

Data used in this work were obtained from the ADNI (http://adni.loni.usc.edu). The ADNI was launched in 2003 as a public-private partnership, led by Principal Investigator Michael W. Weiner, MD. The primary goal has been to test whether serial MRI, PET, other biological markers, and clinical and neuropsychological assessment can be combined to measure the progression of MCI and early AD. For up-to-date information, see (www.adni-info.org).

This study included all ADNI participants with baseline demographics, APOE4, psychological test results, and MRI biomarkers, who also had available longitudinal CSF measurements or 18 F-florbetapir (AV45) PET measurements.

### CSF and PET Measurements

CSF samples were collected and processed as previously described ^13^. The concentration of A*β*42, pTau, and tTau in CSF was measured using the fully automated Elecsys immunoassays (Roche Diagnostics, Basel, Switzerland) by the ADNI biomarker core (University of Pennsylvania, Philadelphia, PA). We obtained these measures from the ADNI depository (UPENNBIOMK9_04_19_17.csv, UPENNBIOMK10_07_29_19.csv, UPENNBIOMK12_01_04_21.csv). CSF A*β*-positivity was defined based on CSF A*β*42 levels with a cutoff value of 880 pg/mL ^14^.

ADNI PET acquisition and processing protocols are described in previously published methods ^15,16^ (https://adni.loni.usc.edu/methods/pet-acquisition/, https://adni.loni.usc.edu/methods/pet-analysis-method/pet-analysis/). We obtained global and regional 18 F-florbetapir SUVR (standardized uptake value ratios) values from the UCBERKELEYAV45_04_26_22.csv table downloaded from the ADNI website (http://adni.loni.usc.edu/). To determine A*β*-positivity, we used a cutoff value of 1.11 to the global SUVR, i.e., summary florbetapir cortical SUV normalized by whole cerebellum SUV ^17^.

### Demographics, APOE4, cognitive measures, and MRI

The ADNI baseline demographics (age, gender, years of education), APOE4, neuropsychological test results, and MRI biomarkers were obtained from ADNIMERGE.csv table downloaded from the ADNI website (http://adni.loni.usc.edu/). We used RAVLT (Reys Auditory Verbal Learning Test) Immediate, RAVLT Learning, RAVLT Forgetting, RAVLT Percent Forgetting, ADAS13 (Alzheimer Disease Assessment Scale-13 items), ADASQ4 (ADAS Delayed Word Recall), MMSE (Mini Mental State Examination), LDELTOTAL (Logical Memory Delayed Recall Total), TRABSCOR (Trail Making Test Part B), FAQ (Functional Assessment Questionnaire), and CDRSB (Clinical Dementia Rating Sum of Boxes) as cognitive measures. These standard measurements, which are widely used in assessing the cognitive and functional performance of dementia patients, are explained in the ADNI General Procedures Manual (http://adni.loni.usc.edu/wp-content/uploads/2010/09/ADNI_GeneralProceduresManual.pdf). As MRI biomarkers, we used the volumetric measures derived from FreeSurfer, which are listed in the ADNIMERGE table. These measures include volumes of ventricles, hippocampus, whole brain, entorhinal, fusiform, middle temporal gyrus, and intra-cranial volume (ICV). However, different quality control measures were considered when constructing MRI biomarkers in the ADNIMERGE table, resulting in a large number of missing values. We decided to omit the quality control criteria and recalculate these MRI biomarkers due to the limited number of data. Omitting the quality control had no significant effect on the results while allowing us to have more data samples.

### Study cohorts

To develop our classification models for predicting progression toward A*β*-positivity, we assigned participants into two overlapping cohorts based on the availability of longitudinal CSF measurements or 18 F-florbetapir (AV45) PET measurements. Specifically, we compiled a CSF-based cohort and a PET-based cohort. The subject selection procedure for each cohort is visualized in Fig. 1.

**Figure 1:**
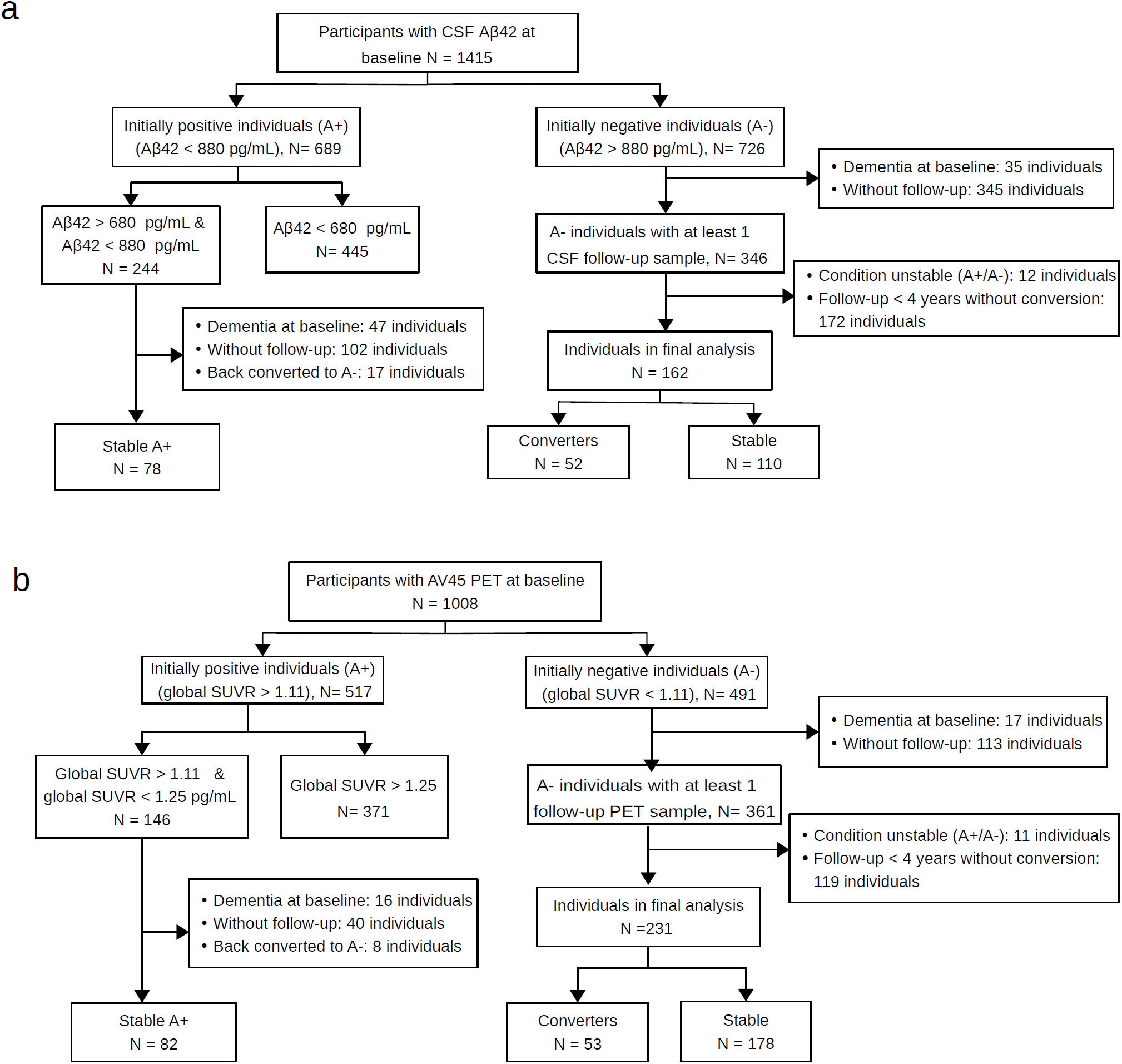
Data selection procedure for predicting progression to A*β*-positivity for a) CSF-cohort, and b) PET-cohort.

For the CSF-based cohort, we selected participants with available baseline CSF data (N= 1415). Among these participants, we first selected the initially A*β*-negative individuals (N = 726) and then we excluded participants 1) with an initial diagnosis of dementia (N= 35), 2) without available follow-up CSF sample (N = 345), 3) with a follow-up duration of fewer than 4 years without conversion to A*β*-positivity (N = 172), and 4) with an unstable A*β*+/A*β*-status during the follow up (N= 12). Finally, among the remaining 162 individuals, we defined A*β*-negative individuals who converted positive within the available follow-up time period as A*β*-Converter (N=52) and individuals who remained A*β*-negative for more than 4 years as A*β*-Stable (N = 110). In order to balance the dataset, we selected a set of auxiliary data from the initially A*β*-positive group. This step was crucial because the number of participants in the A*β*-Converter group was considerably lower than in the A*β*-Stable group, which could negatively impact the classification performance. Specifically, we selected individuals from the initially A*β*-positive group (N= 689) whose A*β*42 values were close to the cutoff used for classifying participants into the A*β*-positive and A*β*-negative groups, i.e., individuals with A*β*42> 680 pg/mL and A*β*42< 880 pg/mL (N= 244). The cut-off points of 680 and 880 were chosen for auxiliary data selection in such a way that a reasonable number of samples were obtained to balance the data set. We further excluded 47 individuals with an initial dementia diagnosis, 102 individuals without a follow-up CSF sample, and 17 individuals who returned to A*β*-negative status during the available follow-up. The remaining participants (N=78) are used as A*β*-Converters, but only during the training phase for balancing the dataset. This approach helped to create a more balanced dataset, which was necessary to develop a robust classification model. The same procedure was applied for selecting the PET-based cohort, which resulted in a dataset of 53 individuals as A*β*-Converters and 178 individuals as A*β*-Stables, with 82 individuals in the auxiliary dataset for use as A*β*-Converters in the training phase.

To develop our regression models for predicting future A*β*42, we only selected subjects with available baseline and longitudinal CSF data. Among those, we eliminated participants with dementia diagnosis at baseline and then we restricted our selection to participants with CSF follow-up samples for at least four years. If an individual had multiple follow-up samples after four years, the first sample (closest to four years) was used as the future sample for the analysis. 253 individuals were included in our dataset as a result. Similarly, we selected 385 individuals for estimating future global SUVR based on PET data. The baseline characteristics of study cohorts are presented in Table 1 and participant’s RIDs are available as supplementary material.

**Table 1:** Characteristics of the study cohorts: Age, education, A*β*42, and global SUVR measures are reported as mean(standard deviation). CN: cognitively normal, SMC: subjective memory concern (participants with self-reported significant memory concern), MCI: mild cognitive impairment, EMCI: early MCI, LMCI: late MCI. Classification of EMCI and LMCI is done by ADNI based on the WMS-R Logical Memory II Story A score. The specific cutoff scores were as follows (out of a maximum score of 25): EMCI was assigned for a score of 9-11 for 16 or more years of education, a score of 5-9 for 8-15 years of education, or a score of 3-6 for 0-7 years of education. LMIC was assigned for a score of 8 for 16 or more years of education, a score of 4 for 8-15 years of education, or a score of 2 for 0-7 years of education ^18^.

| Baseline characteristics | CSF-cohort |  |  |  | PET-cohort |  |  |  |
| --- | --- | --- | --- | --- | --- | --- | --- | --- |
| | Aβ-Converter | Aβ-Stable | Auxiliary data | Regression-cohort | Aβ-Converter | Aβ-Stable | Auxiliary data | Regression-cohort |
| Sample size (N) | 52 | 110 | 78 | 253 | 53 | 178 | 82 | 385 |
| ADNI1/ADNIGO/ADNI2/ADNI3 | 15/6/30/1 | 25/18/67/0 | 27/7/41/3 | 81/35/137/0 | 0/7/44/2 | 0/31/140/7 | 0/8/66/8 | 0/69/308/8 |
| Age, years | 73.7 (7.9) | 71.4 (7) | 73.11 (6.7) | 72.1 (6.7) | 71.9 (6.9) | 70.2 (6.8) | 72.4 (7.1) | 71.4 (6.7) |
| Sex, M/F | 25/27 | 59/51 | 38/40 | 138/115 | 26/27 | 92/86 | 38/44 | 194/191 |
| Education, years | 16.9 (2.4) | 16.3 (2.6) | 16.1 (2.8) | 16.3 (2.7) | 16.6 (2.5) | 16.7 (2.5) | 16.2 (2.7) | 16.4 (2.5) |
| APOE4 (0,1,2) | 32,15,5 | 96,14,0 | 32,36,10 | 163,73,17 | 32,18,3 | 149,26,3 | 40,36,6 | 245,116,24 |
| CN/SMC/EMCI/LMCI | 16/5/12/19 | 51/12/26/21 | 19/6/18/35 | 95/18/62/78 | 28/7/13/5 | 65/24/64/25 | 17/20/28/17 | 123/57/139/66 |
| Aβ42 (CSF-cohort)/global SUVR (PET-cohort) | 1036.4 (182.2) | 1791.4 (526.4) | 754.4 (54.8) | 1196.3 (661.5) | 1.05 (0.04) | 1.004 (0.04) | 1.18 (0.04) | 1.44 (0.20) |

### Machine learning framework

We developed a classifier based on the ridge logistic regression (RLR) approach ^19^ to predict the conversion of A*β*-positivity within 4 years in A*β*-negative individuals in both the CSF-based cohort and PET-based cohort. The framework of the classification procedure is shown in Fig. 2. We designed three models with different feature combinations for each PET-based and CSF-based A*β*-positivity prediction task: The first model was trained using only demographic data and APOE4, the second model was trained using neuropsychological test results in addition to demographic data and APOE4, and the third model was trained using all demographic data, APOE4, neuropsychological test results, and MRI biomarkers. In addition to the classification task, we developed a regression strategy based on ridge linear regression ^20^ to predict future A*β*42 (CSF-based) and global SUVR (PET-based) values from multimodal data similar to the classification task with three models using various feature combinations. By applying a regression model, we were able to eliminate the effects of the cutoff point for classifying participants into A*β*+/A*β*-groups on the performance.

**Figure 2:**
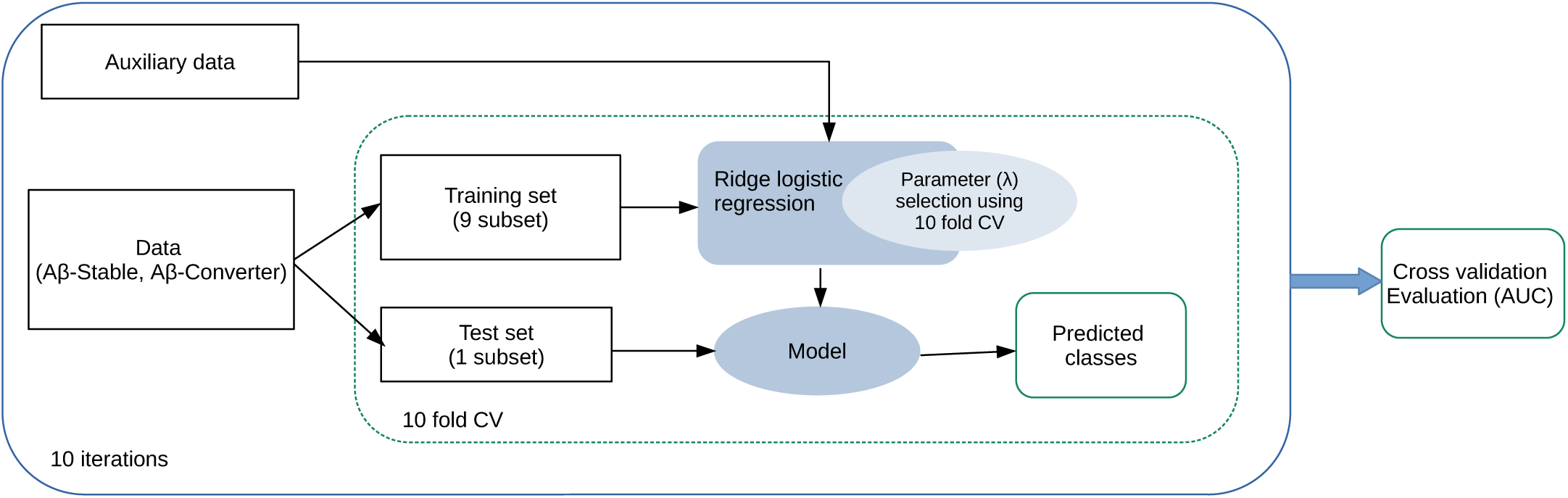
Schematic representation of the classification framework

Moreover, as presented in Table 1, the participants in the CSF-cohort originated from different ADNI cohorts, including ADNI1. The participants in ADNI1 underwent 1.5-Tesla (1.5-T) T1-weighted MRI images, whereas participants in other ADNI cohorts underwent 3T MRI scans. To account for differences in MRI biomarkers caused by different field strengths, we applied ComBat ^21^, as a domain adaptation method to the MRI biomarkers before performing the actual machine learning procedure. ComBat employs an empirical Bayes method for batch correction in microarray expression data, and it has also been successfully utilized for domain adaptation in imaging data ^22–24^. We then compared the results obtained with and without applying ComBat. However, the addition of a domain adaptation step using ComBat approach did not affect the results. Therefore, we excluded it and applied our machine learning approach to the original MRI biomarkers. The codes for classification and regression analyses are available at “https://github.com/ElahehMoradi/AB-positivity-prediction”.

### Implementation and performance evaluation

We used two nested cross-validation loops (10-fold for each loop) to evaluate the model’s performance and estimate the model’s parameter (*λ*) to prevent overfitting and maximize performance. First, an external 10-fold cross-validation was implemented in which samples were randomly divided into 10 subsets with the same proportion of each class label (stratified cross-validation). At each step, a single subset was left for testing and the remaining subsets were used for training. Again, the training set was divided into 10 subsets used to select the models parameter (*λ*). The optimal parameters were selected in classification according to the misclassification error and in regression according to MSE(the mean square error) across the 10-fold of the inner loop. The performance of the model was then evaluated based on mainly AUC (area under the receiver operating characteristic curve) in classification analyses and correlation score in regression analyses in the test subset of the outer loop. We also provided balanced accuracy, sensitivity, and specificity for the classification analyses and mean absolute error for regression tasks in supplementary materials. The reported results in the Results section are averages over 10 nested 10-fold CV runs. Repeated CV was used to reduce variability due to the partitioning of the data.

To compare the AUCs of two learning models, we used the Delong test on the results of a computation run with the median AUC. Comparison of correlation coefficient was tested using methods described by Diedenhofen and colleagues ^25^. All the analyses were done using R (version 4.1.1), with the following pakcages: glmnet ^26^, caret ^27^, sva ^28^, cocor ^29^, pROC ^30^, Daim ^31^, ggplot2^32^, complexheatmap ^33^.

## RESULTS

### Predicting PET and CSF A*β*-positivity in A*β*-negative individuals from multimodal data

We predicted the progression to A*β*-positivity based on CSF and PET data. The number of participants in our PET-cohort and CSF-cohort differs because individuals in each cohort are selected based on the CSF/PET data availability (Fig. 1). Since the PET-based dataset contains more individuals, we subsampled randomly the PET-based dataset to have the same number of individuals in the training phase as the CSF-based model. As explained in the Methods, we designed three models with different feature combinations for each classification of PET-based and CSF-based future A*β*-positivity prediction as well as for the regression model of predicting future A*β*42 and global SUVR values. Fig. 3 shows the results of all these computational analyses. These results are the average over 10 times repeated 10-fold cross-validation analyses for each method.

**Figure 3:**
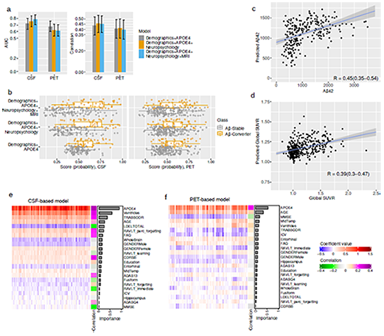
Predicting future A*β*-positivity from multimodal data: a) Bar plots showing the average AUC and average correlation score across 10 computation runs for CSF-based and PET-based models, with 95% confidence intervals error bars. b) Distribution of probability score derived by RLR for PET-based and CSF-based prediction in A*β*-Stable and A*β*-Converter groups. c,d) Scatter plot for estimation of A*β*42 (c) and global SUVR derived by ridge linear regression (with demographics, APOE4, neuropsychology, and MRI biomarkers). The results in b, c, and d are from 1 computation run with median performance e, f) Heatmap of coefficient values across 10 runs of 10-fold CV (100 models) for CSF-based models and pet-based models (f), with a single column heatmap representing the correlation score between each variable and the label (A*β*-Stable, A*β*-Converter), and a bar graph showing the importance of each predictor calculated by the mean of the absolute value of regression coefficients derived by RLR. There are 100 columns in the heatmaps, with each column representing the coefficient values for one model. The performance of predicting A*β*-positivity in A*β*-negative individuals was higher with CSF-cohort compared to PET-cohort, suggesting the higher relevance of CSF data for conversion prediction. ADAS13:Alzheimer Disease Assessment Scale, 13 items, ADASQ4: ADAS Delayed Word Recall, MMSE: Mini-Mental State Examination score, RAVLT: Rey’s Auditory Verbal Learning Test, LDELTOTAL: Logical Memory Delayed Recall Total, TRABSCOR: Trail Making Test Part B, FAQ: Functional Assessment Questionnaire, CDRSB: Clinical Dementia Rating-Sum of Boxes, ICV: intracranial volume, MidTemp: middle temporal gyrus.

According to the findings (Fig. 3 a,b,c,d, Fig. S1), CSF-based models outperformed PET-based models in both classification and regression tasks. The average AUC value across 10 computation runs was 0.78 with a 95% confidence Interval (95% CI) of 0.70 to 0.85 with all demographic, APOE4, neuropsychological test results, and MRI biomarkers for predicting CSF-based progression to A*β*-positivity and 0.61 (95% CI of 0.53 to 0.70) for predicting PET-based progression to A*β*-positivity. The average correlation score for predicting future A*β*42 was 0.45 (95% CI of 0.37 to 0.53), and the average correlation score for predicting future global SUVR was 0.40 (95% CI of 0.31 to 0.49), with all demographic, APOE4, neuropsychological test results, and MRI biomarkers.

The addition of neuropsychological test results and MRI biomarkers to the demographics and APOE4 data increased the performance of the CSF-based A*β*-positivity conversion prediction model as shown in Fig. 3a. The average AUC value for predicting A*β*-positivity in A*β*-negative individuals increased from 0.72 (95% CI of 0.63 to 0.80) to 0.78 (95% CI of 0.70 to 0.85), although the improvement was not statistically significant (p=0.18) according to Delong’s test. The average correlation value for predicting future abeta42 value also increased slightly, from 0.44 (95% CI of 0.35 to 0.52) to 0.45 (p=0.50). However, adding neuropsychological test results and MRI biomarkers to demographic and APOE4 measures did not improve the performance of the PET-based models. Our results showed that including these variables decreased the performance of the PET-based A*β*-positivity conversion prediction model. The average AUC value decreased from 0.68 (95% CI of 0.60 to 0.76) to 0.61 (95% CI of 0.53 to 0.70). The average correlation value for predicting future global SUVR value also slightly decreased, from 0.41 (95% CI of 0.32 to 0.49) to 0.40 (95% CI of 0.31 to 0.49).

We investigated the contribution of different variables in CSF-based and PET-based models, accounting for all demographics, APOE4, neuropsychology test results, and MRI biomarkers (Fig. 3 e & f, Fig. S2). Fig. 3 e & f show the coefficient values for 100 RLR classification models, derived from 10 runs of 10-fold CV, with a single column heatmap representing the correlation score between each variable and the label (A*β*-Stable, A*β*-Converter), and a bar graph indicating the significance of each variable, calculated by taking the mean of the absolute values of the regression coefficients.

In PET-based models, the primary contributors were APOE4 and age (as shown in Fig. 3 f), whereas in CSF-based models (Fig. 3 e), several variables from different data types are significantly contributing, with APOE4 being the most significant, followed by the volume of ventricles and TRABSCORE (Trail Making Test Part B). Additionally, the correlation panels in Fig. 3e & f, and Table S1, and Table S2 show a stronger correlation between various neuropsychological test results and MRI biomarkers with CSF-based labeling compared to PET-based labeling. This explains the higher contribution of different variables in CSF-based modeling, as well as the superior performance of CSF-based modeling.

Moreover, we investigated the correlation between A*β*42 and the A*β*42/A*β*40 ratio with PET global SUVR using ADNI data. The correlation between A*β*42 and PET global SUVR was -0.59, whereas the correlation between the A*β*42/A*β*40 ratio and PET global SUVR measure was -0.73 (Fig. S3). However, we defined A*β*-positivity based on A*β*42 alone, since the A*β*40 measure was only available for a small number of participants.

### Predicting PET and CSF A*β*-positivity in A*β*-negative individuals from PET and CSF baseline measures

We investigated the use of baseline CSF measurements, ie, A*β*42, pTau, and Tau measurements, to predict future CSF-based A*β*-positivity and the use of baseline PET measurements, ie, global and regional SUVR measurements, to predict future PET-based A*β*-positivity. Our goal was to assess the validity of baseline CSF and PET measurements for predicting future conversion. To gain insight into the contribution of other data types besides CSF baseline and PET baseline measurements, we designed four models with different feature combinations. We further developed a regression-based approach to predict the changes (future-baseline) in the A*β*42 and global SUVR values. We predicted the changes in A*β*42/global SUVR values rather than their future values because of the high correlation between the baseline and future values of these measurements. For predicting the changes in A*β*42, we removed 11 individuals from the CSF regression cohort as outliers, with the changes in A*β*42 higher than 500. Similarly, for predicting the changes in global SUVR, we removed 11 individuals from the PET regression cohort as outliers, with the changes in global SUVR less than -0.1. Fig. 4 and Fig. S4 show the results of all these computational analyses. These results are the average over 10 times repeated 10-fold cross-validation analyses for each method.

**Figure 4:**
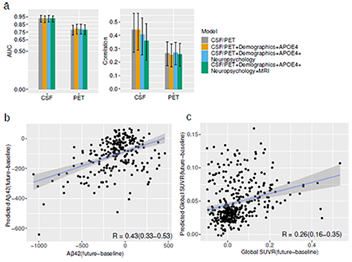
Predicting future A*β*-positivity from baseline CSF/PET measures: a) Bar plots showing the average AUC and average correlation score (predicting the difference between future and baseline A*β*42/global SUVR measures) across 10 computation runs for CSF-based and PET-based models, with 95% confidence intervals error bars, CSF/PET stands for CSF baseline measures (A*β*42, pTau, Tau) for predicting CSF-based A*β*-positivity and PET measures (global and regional) for predicting PET-based A*β*-positivity. b, c) Scatter plot for estimation of the difference between future and baseline A*β*42 (c) and the difference between future and baseline global SUVR (d) derived by ridge linear regression (with CSF/PET measures). The results are from 1 computation run with median performance.

As expected, using baseline CSF measurements to predict future CSF-based A*β*-positivity and using baseline PET measurements for predicting future PET-based A*β*-positivity resulted in relatively high performance. However, CSF-based prediction outperformed PET-based prediction in terms of performance (as shown in Fig. 4a). The average AUC value over 10 computation runs was 0.93 (95% CI of 0.88 to 0.97) for CSF-based prediction, whereas the average AUC value was 0.78 (95% CI of 0.71 to 0.84) for PET-based prediction. Moreover, the average correlation value for predicting the changes in A*β*42 was 0.44 (95% CI of 0.27 to 0.56), whereas for predicting the changes in global SUVR value, the average correlation score was 0.27 (95% CI of 0.17 to 0.35). Interestingly, adding other data types did not improve the performance of classification and regression tasks in both CSF-based and PET-based analyses (Fig. 4a, Fig. S4).

### Predicting CSF-based future A*β*-positivity from CSF and PET baseline measures

We further extended our analysis by predicting CSF-based future A*β*-positivity based on baseline CSF measurements and baseline PET measurements. We selected only individuals with available PET baseline measurements in the CSF-based cohort, resulting in a dataset of 84 A*β*-Stables, 36 A*β*-Converters, and 50 A*β*-positive as auxiliary data (for using as A*β*-Converters only in the training phase to balance dataset). We developed a classification model to predict future A*β*-positivity with three different feature combinations: The first model was trained based on CSF measurements only, the second model based on PET measurements only, and the third model based on both CSF and PET measurements. The results are shown in Fig. 5a and Fig. S5. As anticipated, using baseline CSF measurements for predicting CSF-based future A*β*-positivity resulted in improved performance compared to using baseline PET measurements. Interestingly, the addition of baseline PET measurements with baseline CSF measurements didn’t improve the performance of the classification model for predicting future A*β*-positivity. The average AUC was 0.93 (95% CI of 0.87 to 0.98) using only CSF measurements, 0.82 (95% CI of 0.72 to 0.90) using only PET measurements, and 0.91 (95% CI of 0.85 to 0.96) using both CSF and PET measurements.

**Figure 5:**
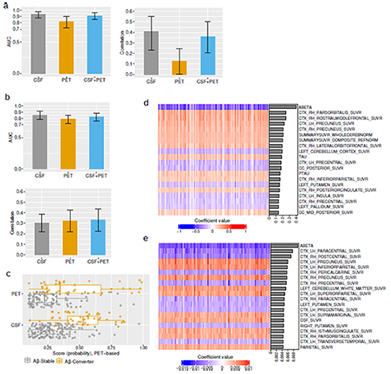
Predicting future CSF-based and PET-based A*β*-positivity from baseline CSF and PET measures: a) Bar plots showing the average AUC and average correlation score across 10 computation runs for predicting future CSF-based amyloid positivity, with 95% confidence intervals error bars. b) Bar plots showing the average AUC and average correlation score across 10 computation runs for predicting future PET-based amyloid positivity, with 95% confidence intervals error bars. c) Distribution of probability score derived by RLR for PET-based prediction in A*β*-Stable and A*β*-Converter groups with CSF and PET baseline measurements. d,e) Heatmap of coefficient values across 10 runs of 10-fold CV (100 models) for PET-based classification model (d) and pet-based regression model (e), with the bar graphs showing the importance of each predictor calculated by the mean of the absolute value of regression coefficient. The regression model is designed for predicting the difference between future and baseline global SUVR measures.

To predict changes in A*β*42, first, we selected 169 individuals from the CSF regression-cohort who had available PET baseline measurements. We then removed six individuals as outliers, with the changes in A*β*42 higher than 500. Again we designed the regression model for predicting the changes in A*β*42 using three different feature combinations: CSF measures only, PET measures only, and the combination of PET and CSF measurements. The resulting average correlation value were 0.41 (95% CI of 0.23 to 0.55) using only CSF measurements, 0.12 (95% CI of 0.004 to 0.25) using only PET measurements, and 0.36 (95% CI of 0.20 to 0.50) using both CSF and PET measurements. Similar to our classification results, the best predictive performance was achieved using only baseline CSF measurements.

### Predicting PET-based future A*β*-positivity from CSF and PET baseline measures

We continued our investigation by predicting PET-based future A*β*-positivity on baseline CSF and PET measurements. Again, we limited the PET-based dataset to individuals with available CSF baseline measurements, resulting in a dataset of 161 A*β*-Stables and 45 A*β*-Converters, and 72 A*β*-positive as auxiliary data (for using as A*β*-Converters only in the training phase to balance dataset). We developed a classification model to predict future PET-based A*β*-positivity with three different feature combinations: The first model was trained based on CSF measurements only, the second model was based on PET measurements only, and the third model was based on both CSF and PET measurements. The results are shown in Fig. 5 b-e and Fig. S6. Surprisingly, the use of baseline CSF measurements to predict future PET-based A*β*-positivity resulted in improved performance compared with the use of baseline PET measurements, the average AUC value increased from 0.79 (95% CI of 0.72 to 0.85) to 0.85 (95% CI of 0.78 to 0.91) (p-value = 0.18). In addition, a combination of CSF and PET measurements provided similar performance to the use of CSF measurements alone, with an average AUC value of 0.83 (95% CI of 0.76 to 0.88). The p-value was 0.049 between the model based on PET measurements alone and the model with the combination of PET and CSF measurements.

To predict changes in global SUVR, first, we selected 353 individuals from the PET regression-cohort who had available CSF baseline measurements. We then removed 10 individuals as outliers, with the changes in global SUVR less than -0.1. Finally, the remaining 343 individuals were used for predicting the changes in global SUVR with different feature combinations. The results are shown in Fig. 5 b and Fig. S6. Unlike the classification results, the average correlation value was rather near to one another in all three different feature combinations. The average correlation value was 0.30 (95% CI of 0.22 to 0.39) with CSF measurements alone, 0.32 (95% CI of 0.22 to 0.42) with PET measurements alone, and 0.33 (95% CI of 0.22 to 0.43) with CSF and PET measurements combined.

We further investigated the contribution of different CSF and PET measurements in both classification and regression models. Fig. 5 d & e show the coefficient values of the 20 variables with the highest importance calculated by taking the mean of the absolute values of the coefficients. Notably, A*β*42 was the most important variable in both classification and regression models (Fig. 5 d & e). Furthermore, the two important CSF measures, i.e., A*β*42 and TAU, rank among the top ten most important variables. These findings demonstrate the importance of CSF measurements in predicting PET-based future A*β*-positivity and also provide an explanation of why the classification model based on baseline CSF measurements outperformed the model based on baseline PET measurements.

### MCI/dementia conversion prediction

Using a completely different data set from the one used for training, we further evaluated our CSF-based and PET-based A*β*-positivity conversion prediction models for MCI/dementia conversion prediction in healthy and MCI individuals. Since A*β* plays a significant role in the initiation of the AD process, the A*β*-positivity conversion prediction model should be able to detect AD-related cognitive decline even though it is not specifically developed to predict conversion to MCI/dementia. To this end, we selected two different datasets for assessing each CSF/PET-based model: The first dataset was for evaluating the future conversion of MCI/dementia in CN individuals and the second dataset was for the evaluation of future conversion of dementia in MCI individuals. For the evaluation of the CSF-based model, we considered all ADNI participants with a baseline diagnosis of CN or MCI who were not utilized for training the CSF-based approach and who had available baseline demographics, APOE4, neuropsychological test results, and MRI biomarkers. We determined two groups based on baseline and longitudinal diagnosis labels, regardless of CSF/PET biomarker status: a) Stable group: CN/MCI individuals who have remained stable for five years or more following baseline, b) Converter group: 1) Individuals with CN diagnosis at baseline who later develop MCI or dementia (the last two diagnoses must be MCI or dementia), and 2) MCI individuals who later develop to dementia (the last two diagnoses must be dementia). This resulted in a dataset of 67 converters and 131 stables for conversion prediction in CN individuals and a dataset of 211 converters and 126 stables in conversion prediction in MCI individuals (Table S3). The same procedure was used to select data for the PET-based model evaluation, resulting in a dataset of 74 converters and 95 stables for conversion prediction in CN individuals and a dataset of 234 converters and 90 stables in conversion prediction in MCI individuals (Table S4).

We trained a model for predicting CSF-based future A*β*-positivity using all available subjects (Fig. 1 a, Table 1) and used it to classify CN-Stable vs. CN-Converter groups, as well as MCI-Stable vs. MCI-Converter groups. We next repeated the process for the PET-based model. The results are shown in Fig. 6. The CSF-based model performed quite well, achieving an AUC of 0.76 (95% CI of 0.68 to 0.83) for MCI/dementia conversion prediction in the CN group and an AUC of 0.89 (95% CI of 0.86 to 0.93) for dementia conversion prediction in the MCI group. However, the PET-based model had lower performance, with an AUC of 0.60 (95% CI of 0.51 to 0.69) for classifying CN-Stables vs. CN-Converters and an AUC of 0.72 (95% CI of 0.67 to 0.78) for classifying MCI-Stables vs. MCI-Converters.

**Figure 6:**
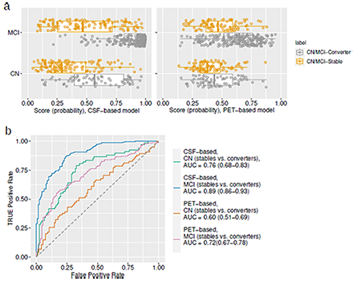
MCI/dementia conversion prediction in CN and MCI groups: a) Boxplot of probability score derived by RLR for CSF-based and PET-based model for classification of stables vs converters in CN and MCI groups. b) ROC curves of CN and MCI subjects classification to stable and converter groups using CSF-based and PET-based models, with AUC values and 95% confidence intervals.

Given that neither the CSF-based nor the PET-based model was specifically designed for classifying CN-Stables vs. CN-Converters/MCI-stables vs. MCI-converters, and that both models used basic and noninvasive data (demographics, APOE4, cognitive, and MRI), the CSF-based model performed exceptionally well for the MCI/dementia conversion prediction in CN and MCI groups, while the PET-based model performed well only in the MCI group. However, in the CN group, the PET-based model failed to predict conversion to MCI/dementia.

## DISCUSSION

Determining A*β*-status is crucial for the prescription of amyloid-targeted treatments in the future. This study aimed to predict conversion to A*β*-positivity in A*β*-negative individuals using data that is widely available in clinical settings. To achieve this goal, we developed a classification model based on an RLR approach incorporating demographics, APOE4, neuropsychological tests, and MRI biomarkers. We categorized participants into A*β*-positive and A*β*-negative groups, based on CSF A*β*42 and PET global SUVR. Due to categorization inconsistencies ^34^, we conducted separate analyses with participants classified as A*β*+/A*β-*based on CSF and PET biomarkers. Additionally, we predicted future CSF A*β* (A*β*42) or PET A*β* (global SUVR) values via a ridge linear regression approach to eliminate any bias arising from the cutoff point for categorizing participants into A*β*+/A*β*-groups on the performance.

In our analyses, we demonstrated that features from multiple modalities, including demographics, neurophysiological scores, APOE4, and MRI biomarkers, can model the progression to A*β*-positivity detected by either CSF or PET biomarkers. Interestingly, our findings indicate that utilizing CSF A*β*42 for participant categorization resulted in more accurate predictions of future A*β*-positivity compared to PET global SUVR. Also, the CSF-based model benefited from additional features, whereas the accuracy of the PET-based model decreased with more features. In more detail, the AUC of CSF-based model increased from 0.72 with APOE and demographics as the features to 0.78 with all the features. However, the AUC of the PET-based decreased from 0.67 with APOE and demographics as the features to 0.61 to all the features. To comprehend the performance disparity between the PET-based and CSF-based models, we investigated the contributions of different data types in each model. We observed that cognitive scores and MRI biomarkers, along with APOE4 as the most important variable, strongly contribute to the CSF-based model. In contrast, in the PET-based model, the contribution of cognitive and MRI biomarkers was smaller, and the primary predictors were APOE4 and age.

It is essential to highlight that CSF A*β*42 and PET global SUVR measures, utilized in defining A*β*-positivity, employ different mechanisms for detecting A*β* protein. PET imaging reveals the presence of amyloid plaques in the brain, whereas CSF analysis is associated with the clearance of amyloid from the brain. Low CSF amyloid levels may indicate inefficient clearance, leading to brain amyloid accumulation ^35,36^. Although we used the same ML framework for predicting future A*β*-positivity, direct comparison of the results is challenging. However, our findings suggest that predicting future CSF-based A*β*-positivity is relatively easier compared to PET-based A*β*-positivity.

We defined A*β*-positivity based on A*β*42 alone, since the A*β*40 measure was only available for a small number of participants. However, recent studies emphasize the relevance of the A*β*42/A*β*40 ratio in defining A*β*-positivity using CSF biomarkers ^37^ as it correlates more strongly with PET-based A*β* measures compared to A*β*42 alone. We speculate that utilizing the A*β*42/A*β*40 ratio for predicting CSF-based A*β*-positivity might yield results that are more aligned with PET-based predictions. However, due to the limited number of available samples, we cannot conduct experiments to confirm this hypothesis.

Previous studies have primarily focused on detecting A*β*-positivity at the time of the study ^38–42^ (baseline prediction). These studies have employed either machine learning algorithms or statistical analyses to identify A*β*-positivity at baseline and explore the relationships between cognitive measures, various biomarkers, and A*β*-positivity. For example, Palmqvist et al. ^38^ developed a model utilizing demographics, cognitive tests, white matter lesions, APOE, and plasma biomarkers (A*β*42/A*β*40, tau, and neurofilament light chain) to detect A*β*-positivity at baseline. Their model achieved an AUC of 0.80-0.82 when trained on BIOFINDER dataset ^39,43^ and validated on ADNI data. Similarly, another study by the same group ^39^, detected baseline A*β*-positivity using demographic, APOE, and cognitive information and achieved an AUC of 0.65 in cognitively healthy individuals. However, we are aware of only two studies that have focused on predicting future A*β*-positivity ^12,44^. The first study by Elman et al. ^44^ examined the association of baseline cognitive measures with progression to A*β*-positivity in A*β*-negative individuals. The second study conducted by Park et al. ^12^ employed machine learning algorithms to predict future A*β*-positivity in A*β*-negative individuals. In Park et al. ^12^ study, they used a PET biomarker for subject classification in A*β*+/A*β*-groups and developed a classifier with baseline age, gender, APOE4 genotype, and PET SUVR measures in ADNI data. They achieved a cross-validated AUC of 0.67 using basic demographic and genetic factors, which improved to 0.84 when including PET SUVR measures. Our work has a similar objective to the study by Park et al. ^12^, but we focused on widely available non-invasive measures to predict future A*β* conversion. Moreover, a key characteristic of our study was to assess the future predictability of A*β*-positivity determined based on either CSF and PET biomarkers whereas Park et al. ^12^ only considered a PET-based definition.

We validated the relevance of our prediction models for future A*β*-positivity in an independent dataset composed of separate ADNI participants, predicting clinical status changes from CN to MCI/dementia or from MCI to dementia. The CSF-based model performed well, achieving an AUC of 0.76 for MCI/dementia conversion prediction in CN individuals and an AUC of 0.89 for the dementia conversion prediction in MCI individuals. However, the PET-based model performed worse in predicting conversion, reaching an AUC of 0.60 for CN to MCI/dementia conversion and an AUC of 0.72 for MCI to dementia conversion. The CSF-based model’s excellent performance in predicting MCI/dementia conversion in a completely independent dataset is intriguing, indicating its potential for detecting AD/dementia-related changes at an early stage. Although the model was not specifically designed to classify CN/MCI-Converter versus CN/MCI-Stable groups, the CSF-based model demonstrated strong performance comparable to existing studies designed for such classification tasks ^45–47^.

We analyzed the use of baseline CSF and PET measurements for predicting future A*β*-positivity. Including baseline scores significantly improved the performance of the prediction models. Notably, when baseline CSF/PET measurements were available, the addition of other data types such as cognitive scores and MRI biomarkers contributed very little value. Moreover, utilizing baseline CSF/PET measurements enabled the prediction of the change/rate in A*β*42 and global SUVR values which are rather hard to model ^48^. However, the invasive nature, high cost, and limited availability of CSF/PET measurements restrict their usage. Similarly, we investigated the role of baseline CSF and PET measurements in both predicting CSF-based future A*β*-positivity as well as in predicting PET-based future A*β*-positivity. Interestingly, baseline CSF measurements showed superior predictive ability for future PET-based A*β*-positivity compared to baseline PET measurements. A closer look at the coefficient values in the PET-based model revealed that A*β*42 made the greatest contribution to the model. Recent studies suggest that AD-related alterations are detectable earlier in CSF than in PET ^49,50^, which may explain the superior performance of baseline CSF measurements in predicting PET-based A*β*-positivity.

Our study has several limitations. First, the sample sizes were small due to our specific inclusion criteria, although the data came from one of the largest longitudinal dementia-prediction cohorts (ADNI). To compensate for small sample size, we evaluated our CSF-based/PET-based models in an independent dataset for MCI/dementia conversion prediction. Second, the sizes of A*β*-Stable and A*β*-Converter groups were different, which we partially addressed by introducing auxiliary data for label balancing during model training. Third, the CSF-based and PET-based analyses used different datasets, challenging direct performance comparison. To address this, we included an equal number of participants for CSF-based and PET-based model training to facilitate a fairer comparison. Last, we utilized a single ML approach to combine various data types and develop our models. While this approach worked well for CSF-based analyses, a more advanced approach may be needed for PET-based future A*β*-positivity prediction.

In conclusion, we developed ML-based models to predict future A*β*-positivity in A*β*-negative individuals, determined based on both CSF and PET biomarkers. Higher predictive performance was achieved with CSF-derived dichotomization (AUC = 0.78) compared to PET dichotomization(AUC = 0.68). The discrepancy in performance may be attributed to the different mechanisms of A*β* detection and the discordance between the two biomarkers. Further research is required to understand this discrepancy. However, by using non-invasive measures including demographics, APOE4, neuropsychological scores, and MRI biomarkers, our CSF-based A*β*-positivity conversion prediction model performed well in identifying A*β*-Stables vs A*β*-converters with an AUC 0f 0.78, as well as in identifying CN-Converters vs. CN-Stables (AUC = 0.76) and MCI-Stables vs. MCI-Converters (AUC = 0.89). These findings demonstrate the significance of neuropsychological and MRI biomarkers to detect the risk of conversion to AD, even in cognitively normal individuals. The detection may occur before current thresholds for A*β*-positivity are reached, providing an opportunity for early intervention.

## ACKNOWLEDGMENTS

This research has been supported by European Research Council (grant 804371), The Academy of Finland (grants 346934 and 349184), and a grant 351849 from the Academy of Finland under the frame of ERA PerMed (”Pattern-Cog”), a grant AC21-2/00023 from Spain’s Instituto de Salud Carlos III with NextGenEU European funding from the “Plan de Recuperación, Transformación y Resiliencia (PRTR)”.

Data collection and sharing for this project was funded by the Alzheimer’s Disease Neuroimaging Initiative (ADNI) (National Institutes of Health Grant U01 AG024904) and DOD ADNI (Department of Defense award number W81XWH-12-2-0012). ADNI is funded by the National Institute on Aging, the National Institute of Biomedical Imaging and Bioengineering, and through generous contributions from the following: AbbVie, Alzheimers Association; Alzheimers Drug Discovery Foundation; Araclon Biotech; BioClinica, Inc.; Biogen; Bristol-Myers Squibb Company; CereSpir, Inc.; Cogstate; Eisai Inc.; Elan Pharmaceuticals, Inc.; Eli Lilly and Company; EuroImmun; F. Hoffmann-La Roche Ltd and its affiliated company Genentech, Inc.; Fujirebio; GE Healthcare; IXICO Ltd.; Janssen Alzheimer Immunotherapy Research & Development, LLC.; Johnson & Johnson Pharmaceutical Research & Development LLC.; Lumosity; Lundbeck; Merck & Co., Inc.; Meso Scale Diagnostics, LLC.; NeuroRx Research; Neurotrack Technologies; Novartis Pharmaceuticals Corporation; Pfizer Inc.; Piramal Imaging; Servier; Takeda Pharmaceutical Company; and Transition Therapeutics. The Canadian Institutes of Health Research is providing funds to support ADNI clinical sites in Canada. Private sector contributions are facilitated by the Foundation for the National Institutes of Health (www.fnih.org). The grantee organization is the Northern California Institute for Research and Education, and the study is coordinated by the Alzheimers Therapeutic Research Institute at the University of Southern California. ADNI data are disseminated by the Laboratory for Neuro Imaging at the University of Southern California.

