## Supplementary materials for "Machine learning prediction of future amyloid beta positivity in amyloid-negative individuals"

June 29, 2023

#### Spplementary Tables

Table S1: The correlation between cognitive measures to the labels in CSF-cohort and PET-cohort (A$\beta$-Stable vs. A$\beta$-Converter, and to the labels in regression cohort of CSF (A$\beta$42) and PET(global SUVR).

| | CDRSB | ADAS13 | ADASQ4 | MMSE | RAVLT-immediate | RAVLT-learning | RAVLT-forgetting | RAVLT_perc-forgetting | LDELTOTAL | TRABSCORE | FAQ |
|---|---|---|---|---|---|---|---|---|---|---|---|
| **CSF-cohort** | 0.24 | 0.18 | 0.15 | -0.22 | -0.19 | -0.12 | 0.11 | 0.20 | -0.21 | 0.21 | 0.20 |
| **PET-cohort** | -0.1 | -0.06 | -0.08 | -0.13 | -0.03 | -0.01 | 0.03 | 0.002 | 0.04 | 0.06 | -0.06 |
| **CSF, Regression-cohort** | -0.26 | -0.30 | -0.29 | 0.19 | 0.23 | 0.13 | -0.17 | -0.27 | 0.29 | -0.18 | -0.22 |
| **PET, Regression-cohort** | 0.22 | 0.18 | 0.16 | -0.13 | -0.18 | -0.06 | 0.13 | 0.18 | -0.23 | 0.22 | 0.22 |

Table S2: The correlation between MRI measures (volumes) to the labels in CSF-cohort and PET-cohort (A$\beta$-Stable vs. A$\beta$-Converter, and to the labels in regression cohort of CSF (A$\beta$42) and PET(global SUVR).

| | ICV | hippocampus | Entorhinal | Fusiform | MidTemp | Ventricles | WholeBrain |
|---|---|---|---|---|---|---|---|
| **CSF-cohort** | 0.04 | -0.05 | -0.11 | -0.15 | -0.06 | 0.25 | -0.09 |
| **PET-cohort** | 0.02 | -0.05 | 0.06 | 0.02 | 0.05 | 0.1 | 0.02 |
| **CSF, Regression-cohort** | -0.07 | 0.16 | 0.12 | 0.09 | 0.06 | -0.19 | 0.01 |
| **PET, Regression-cohort** | 0.03 | -0.17 | -0.1 | -0.02 | -0.05 | 0.12 | -0.04 |

Table S3: Characteristics of the Validation cohorts for performance validation of CSF-based model.

| | CN-cohort | | MCI-cohort | |
|---|---|---|---|---|
| **Baseline characteristics** | Converter-CN | Stable-CN | Converter-MCI | Stable-MCI |
| Sample size (N) | 67 | 131 | 211 | 126 |
| ADNI1/ADNIGO/ ADNI2/ADNI3 | 32/0/31/4 | 57/0/73/1 | 145/9/48/19 | 41/40/45/0 |
| Age, years | 76.2 (4.96) | 72.9 (5.4) | 74 (6.7) | 70.7 (7.1) |
| Sex, M/F | 41/26 | 58/73 | 129/82 | 78/48 |
| Education, years | 16.3(2.7) | 16.4 (2.8) | 15.8 (2.8) | 16.0 (2.3) |
| APOE4 (0/1/2) | 42/23/2 | 101/30/0 | 68/105/38 | 84/36/6 |
| CN/SMC/EMCI/LMCI | 51/16/0/0 | 105/26/0/0 | 0/0/24/187 | 0/0/74/52 |

Table S4: Characteristics of the Validation cohorts for performance validation of PET-based model.

| | CN-cohort | | MCI-cohort | |
|---|---|---|---|---|
| **Baseline characteristics** | Converter-CN | Stable-CN | Converter-MCI | Stable-MCI |
| Sample size (N) | 74 | 95 | 234 | 90 |
| ADNI1/ADNIGO/ ADNI2/ADNI3 | 48/0/22/4 | 69/0/24/2 | 165/8/53/8 | 47/25/18/0 |
| Age, years | 75.9 (4.5) | 724.6 (5.6) | 74.1 (6.6) | 72.5 (7.3) |
| Sex, M/F | 46/28 | 42/53 | 147/87 | 34/56 |
| Education, years | 16.1(2.8) | 16.5 (2.8) | 15.9 (2.8) | 15.7 (2.97) |
| APOE4 (0/1/2) | 48/24/2 | 75/19/1 | 80/112/42 | 53/33/4 |
| CN/SMC/EMCI/LMCI | 63/11/0/0 | 80/15/0/0 | 0/0/24/187 | 0/0/22/212 |

### Spplementary Figures

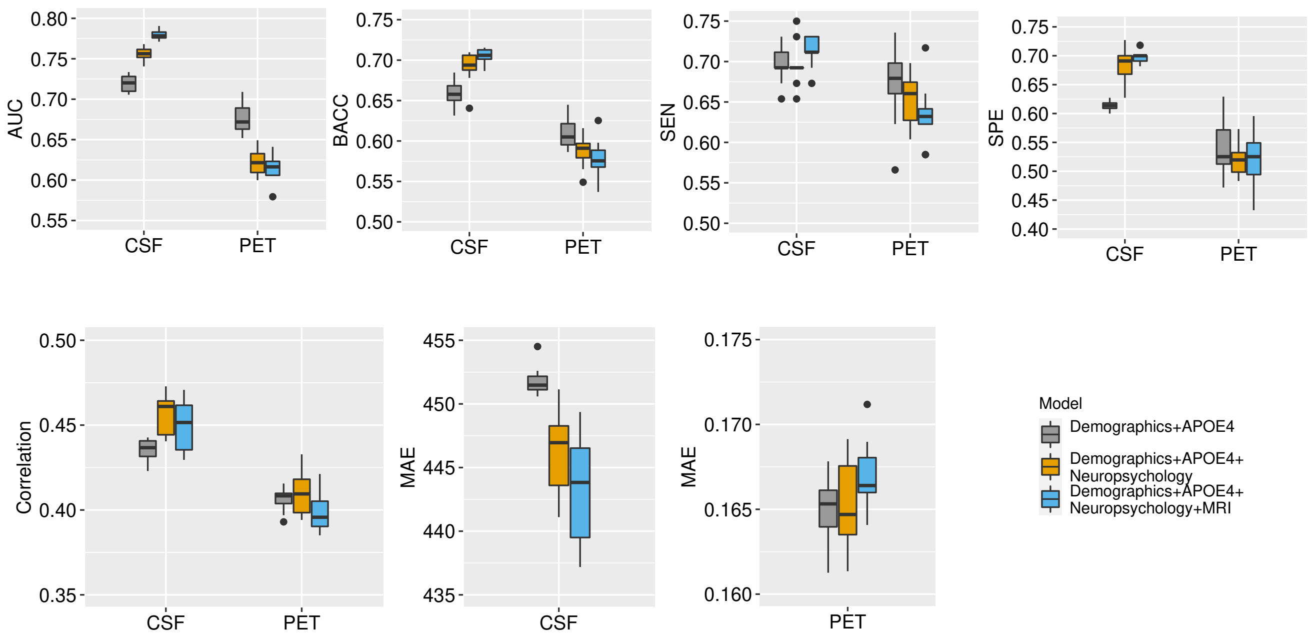


Figure S1: Predicting future A$\beta$ positivity from multimodal data: Box plots for AUC, balanced accuracy (BACC), sensitivity (SEN), and specificity (SPE) for predicting A$\beta$ positivity in A$\beta$ negative individuals, and the correlation score and mean absolute error (MAE) for predicting future A$\beta$42 (CSF) and global SUVR (PET) measures in CSF and PET cohorts. The results are derived from 10 computation runs. In each box, the central mark shown in black is the median, the edges of the box are the 25th and 75th percentiles, whereas the whiskers extend to the most extreme data points not considered outliers, and outliers are plotted as dots.

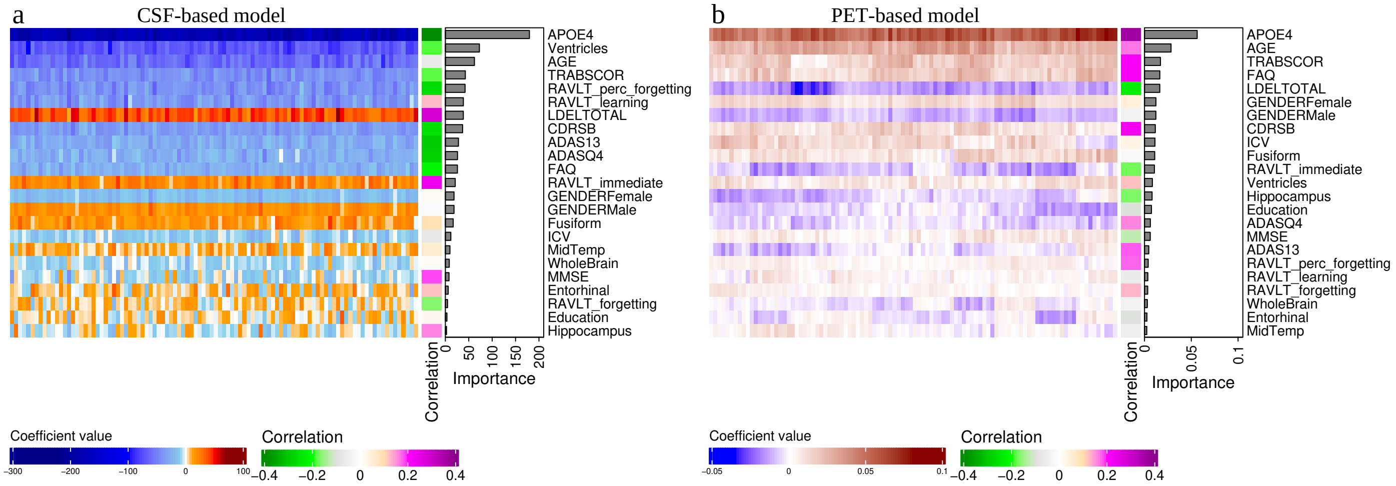


Figure S2: Predicting future A$\beta$ positivity from multimodal data: Heatmap of coefficient values across 10 runs of 10-fold CV (100 models) for a) predicting future A$\beta$42 (CSF-based) and for (b predicting future global SUVR measure (pet-based), with a single column heatmap representing the correlation score between each variable and the label (future A$\beta$42, future global SUVR), and a bar graph showing the importance of each predictor calculated by the mean of the absolute value of regression coefficients derived by ridge linear regression.

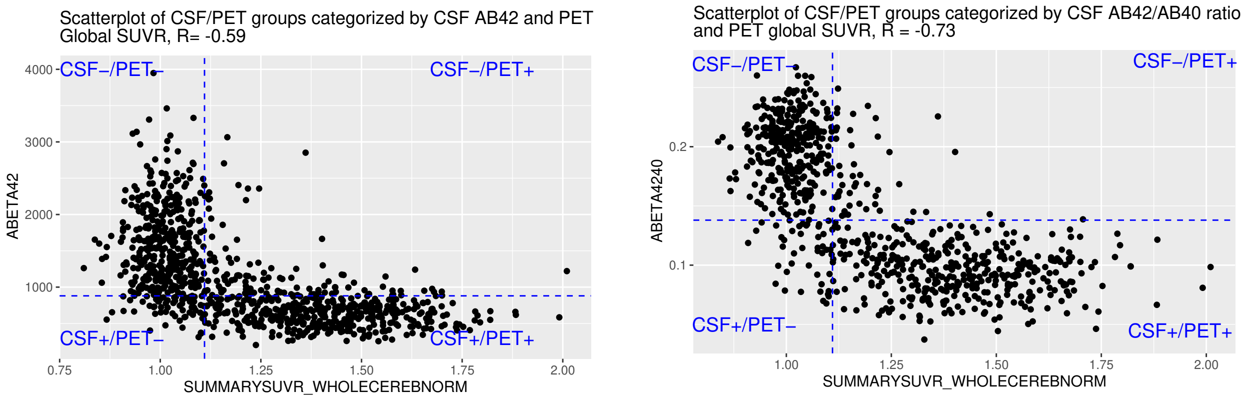


Figure S3: Scatterplots of CSF A$\beta$42 and the A$\beta$42/A$\beta$40 ratio and PET global SUVR.

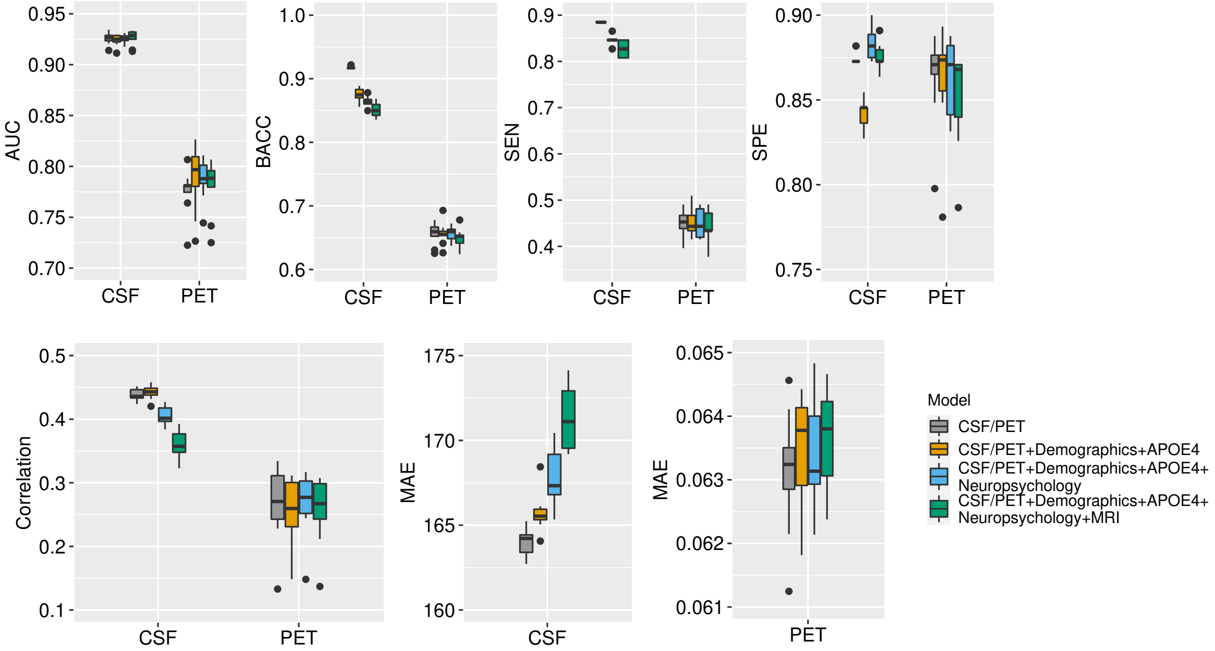


Figure S4: Predicting future A$\beta$ positivity from baseline CSF/PET measures: Box plots for AUC, BACC, SEN, and SPE for predicting A$\beta$ positivity in A$\beta$ negative individuals, and the correlation score and mean absolute error (MAE) for predicting the difference between future and baseline A$\beta$42 (CSF) and global SUVR (PET) in CSF and PET cohorts. The results are derived from 10 computation runs. In each box, the central mark shown in black is the median, the edges of the box are the 25th and 75th percentiles, whereas the whiskers extend to the most extreme data points not considered outliers, and outliers are plotted as dots. CSF/PET stands for CSF baseline measures (A$\beta$42, pTau, Tau) for predicting CSF-based A$\beta$ positivity and PET measures (global and regional) for predicting PET-based A$\beta$ positivity

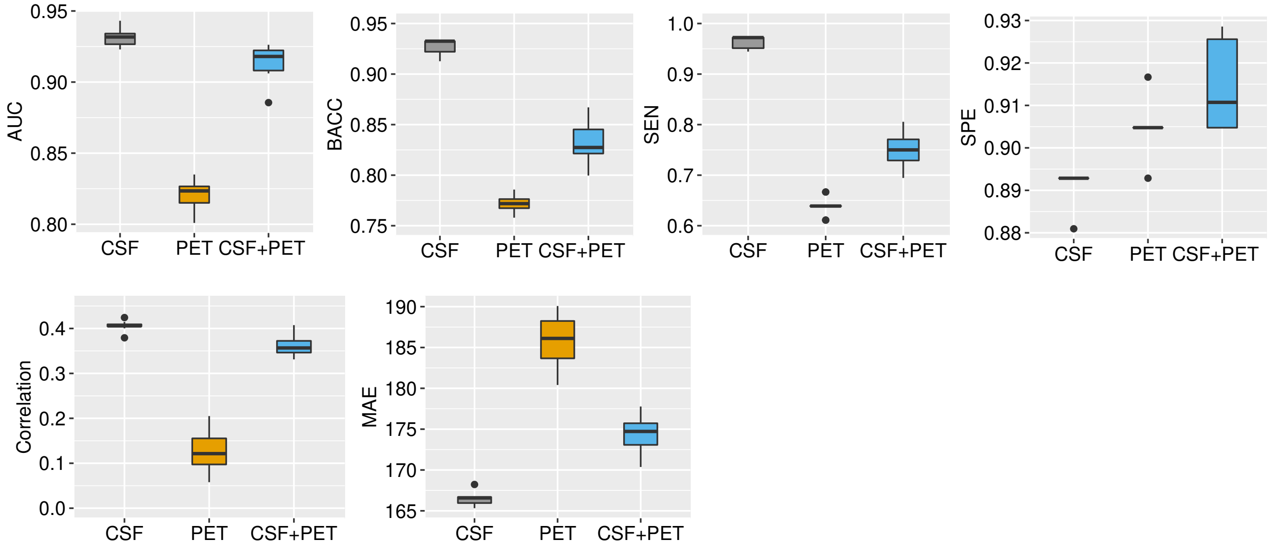


Figure S5: Predicting CSF-based future A$\beta$ positivity from CSF and PET baseline measures: Box plots for AUC, balanced accuracy (BACC), sensitivity (SEN), and specificity (SPE) for predicting A$\beta$ positivity in A$\beta$ negative individuals and the correlation score and mean absolute error (MAE) for predicting the difference between future and baseline A$\beta$42 (CSF-based). The results are derived from 10 computation runs. In each box, the central mark shown in black is the median, the edges of the box are the 25th and 75th percentiles, whereas the whiskers extend to the most extreme data points not considered outliers, and outliers are plotted as dots. CSF stands for CSF baseline measures (A$\beta$42, pTau, Tau) and PET stands for PET baseline measures (global and regional).

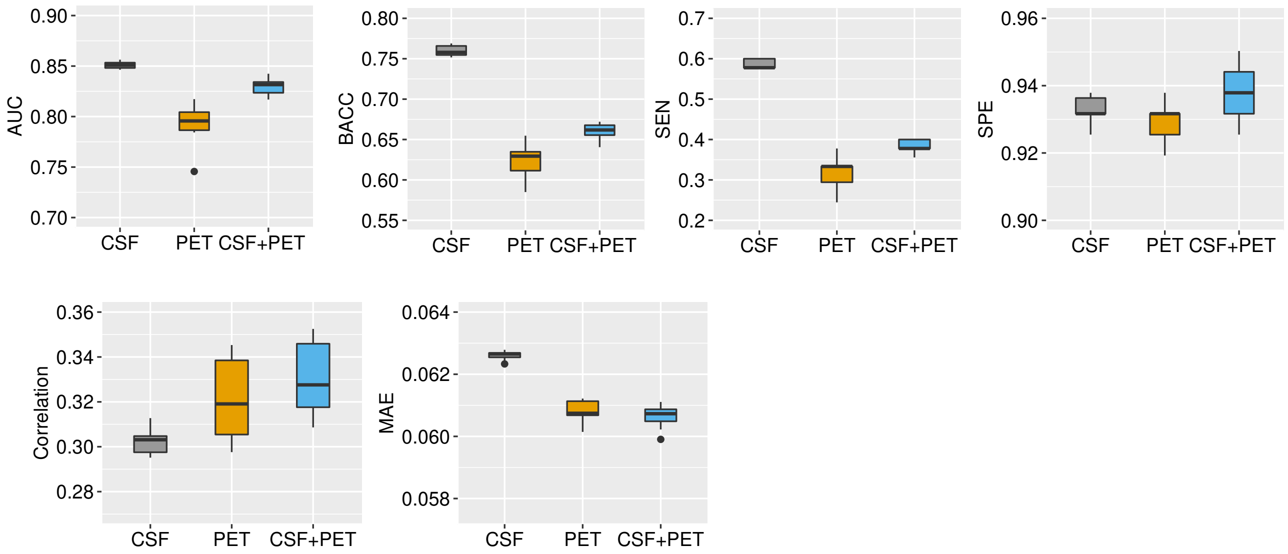


Figure S6: Predicting PET-based future A$\beta$ positivity from CSF and PET baseline measures: Box plots for AUC, balanced accuracy (BACC), sensitivity (SEN), and specificity (SPE) for predicting A$\beta$ positivity in A$\beta$ negative individuals and the correlation score and mean absolute error (MAE) for predicting the difference between future and baseline global SUVR (PET-based). The results are derived from 10 computation runs. In each box, the central mark shown in black is the median, the edges of the box are the 25th and 75th percentiles, whereas the whiskers extend to the most extreme data points not considered outliers, and outliers are plotted as dots. CSF stands for CSF baseline measures (A$\beta$42, pTau, Tau) and PET stands for PET baseline measures (global and regional).
